# Fair allocation of healthcare research funds by the European Union?

**DOI:** 10.1101/453209

**Authors:** Zoltán Kaló, Loek Hendrik Matheo van den Akker, Zoltán Vokó, Marcell Csanádi, György János Pitter

## Abstract

This study aimed to investigate the distribution of European Union (EU) healthcare research grants across EU countries, and to study the effect of the potential influencing factors on grant allocation. We analysed publicly available data on healthcare research grants from the 7th Framework Programme and the Horizon 2020 Programme allocated to beneficiaries between 2007 and 2016. Grant allocation was analysed at the beneficiary-, country-, and country group-level (EU-15 versus newer Member States, defined as EU-13). The investigated country-level explanatory variables included GDP per capita, population size, overall disease burden, and healthcare research excellence. Grant amounts per 100,000 inhabitants was used as an outcome variable in the regression analyses.

Research funds were disproportionally allocated to EU-15 versus the EU-13, as 96.9% of total healthcare grants were assigned to EU-15 countries. At the beneficiary level, EU funding was positively influenced by participating in previous grants. The average grant amount per beneficiary was higher for EU-15 organizations. In univariate regression analyses at the country level, higher GDP per capita (p<0.001) and better medical research excellence (p<0.001) were associated with more EU funding, and a higher disease burden was associated with less EU funding (p=0.003). In the multiple regression analysis GDP per capita (p=0.002) and research excellence (p<0.001) had a significant positive association with EU funding. Population size had an inverted U-shaped relationship with EU funding for healthcare research, having the largest per capita funding in second and the third quartiles (p=0.03 and p=0.02).

The uneven allocation of healthcare research funds across EU countries was influenced by GDP per capita, medical research excellence and population size. Wealthier countries with an average population size and strong research excellence in healthcare had more EU funding for healthcare research. Higher disease burden apparently was not associated with more EU research funding.

## Introduction

Funding is one of the main drivers of scientific activities.^1^ It is important for defining new scientific research projects or improving research and development in existing projects. Research and development is a major field for innovation, employment and economic stimulation in the European Union (EU). Since this field is important to the worldwide economic positioning of the EU, there is considerable funding available to financially support companies, universities and research authorities.^2^

Health contributes signficantly to the economy of the European Union^3^, and health systems are fundamental parts of Europe’s social infrastructure.^4^ Consequently, a shared commitment to health can help to make the value of Europe real for its citizens, and to keep the European economy sustainable and competitive in the coming decades.^5^ Due to differences in health outcomes, quality and cost of care, European research on health systems has great potential to support countries in improving the outcomes and efficiency of their healthcare systems.^6^ Since different countries follow various research patterns, and their economic and institutional structures already greatly differ,^7^ the allocation of European healthcare research funds among countries has the potential to influence current inequalities among member states.

Currently there is limited information about the allocation patterns of research funds and the factors that may influence their distribution across countries. However, successful proposals for research grants could be associated with several features of the applicant. A previous study showed that research centres that are already members of large scientific teams and those who develop a good connection with productive researchers and ensure the flow of information, have higher chances of securing more research funding.^8^ Other studies in the field of healthcare showed that more funding was allocated for illnesses with greater disease burden^9^ and there is an association between receiving research funds and the quality of published medical education research.^10^

Restricted access to research grants predisposes lower research performance, which may eventually further reduce the success rate of research proposals for funding. If funding for research is limited in a country, the scientific ranking and performance of researchers decrease compared to other countries. On a macro scale, this may result in the lower-level development of economic, healthcare and scientific performance and increases the brain drain of researchers from countries with poor research performance.

Several country-specific factors may influence grant allocation at an international level, including the (1) economic status, (2) population size, (3) disease burden and (4) excellence of scientific research capabilities. When considering equity principles and fair distribution, grant allocation across countries should not depend on the economic status of a country and its population size (i.e. greater negotiation power) but should depend on the need for healthcare research because of a higher disease burden and on the excellence of scientific research capabilities.

This research aimed to examine the distribution of EU grants for healthcare among EU member countries and to study the effect of potential influencing factors on grant allocation.

## Methods

### Setting

The time scale of this study was from the starting date of the 7th Framework Programme (FP7) until 31 December 2016. The assigned healthcare grants of the FP7 and Horizon 2020 programmes were exported from the EU Cordis database.^11^

### Data sources

7th Framework Programme: Before the introduction of the current EU funding research programme Horizon 2020 in 2014, the EU funding programme was called FP7. FP7 funds were available for research purposes and contained five project groups, namely, Cooperation, Ideas, People, Capacities and Nuclear Research. These five groups were also divided into different subgroups based on the research subject.^12^ The core of FP7, which represented two-thirds of the overall budget, was the Cooperation programme. It fostered collaborative research across European and other partner countries through projects by transnational consortia of industry and academia. The Cooperation programme contained ten subgroups, including the health subgroup, which is the subject of this study.^13^

Horizon 2020: The EU’s Horizon 2020 funding programme is the EU’s largest research and innovation funding programme since the EU was established, containing nearly 80 billion EUR for a seven-year timeframe (2014-2020). This new programme of the EU aims at stimulating innovation and securing Europe’s position in global competitiveness.^14^ The Horizon 2020 funding programme is available for general research purposes. The three main pillars of the project are excellent science, industrial leadership and social challenges. These three pillars are divided into different subgroups based on the research subject. The EU has identified seven priority challenges wherein targeted research and innovation investments can benefit citizens. The social challenges pillar contains the health subgroup (Horizon 2020-EU-3.1), which is the interest of this study.^14^

### Data collection

There were 1,008 health grants included from the FP7-Health programme and 544 from Horizon 2020. Since Horizon 2020 was still ongoing during this study, the 544 included grants were just a subset of the total funding in the programme. Thus, a total of 1,552 health grants were included in this analysis.

The basic information collected on the included grants contained the grant IDs, the total grant value of the participating beneficiaries, and the grants’ timescale in addition to an overview of the participating beneficiaries. For the purpose of this study, first, the participating beneficiaries were extracted to multiple rows in Microsoft Excel with an R-script. Second, the grant amounts per beneficiary, the corresponding country and the role of the beneficiary (e.g. project coordinator) were manually matched with the Cordis database.^11^ Additional general grant information (i.e., the start date, end date, grant acronym, topic, title and objective) was matched from the general grant programme that was exported by the grant ID number. All grants were labelled with the year of assignation and grant amounts of beneficiaries were mapped to the corresponding countries manually.

The final dataset contained 14,446 participating beneficiaries for the 1,552 assigned grants. The beneficiaries could have participated in multiple grants, therefore the number of unique beneficiaries was lower. For all beneficiaries appearing in more than one grant, their first participation was determined, and grant amounts were calculated for the first and the subsequent grants separately. The dataset contained values for beneficiaries worldwide. However, the focus of this research was the grant distribution among the EU countries only. After the exclusion of beneficiaries from non-EU countries, the final dataset contained 1529 grants with 12,678 EU-beneficiaries.

### Descriptive analysis of the grant distribution

The sum of total assigned grant amounts, and the total assigned grant amounts per 100,000 inhabitants were calculated for all beneficiaries and for all EU countries based on the location of the beneficiaries. The normalisation for the population was necessary to make the outcome comparable among countries with various population sizes. In addition to the institutional and country-level analyses, the grant distribution across EU-15 (member states in the EU prior to the accession of ten candidate countries on 1 May 2004)^15^ and EU-13 countries (member states in the EU joined after 1 May 2004)^16^ was calculated. The frequency of participation and average funding of EU-15 beneficiaries and EU-13 beneficiaries were calculated separately.

The following additional descriptive statistical calculations were conducted at a country group-level with the mean, the median and the interquartile range (IQR) of assigned healthcare grants. The grant amounts were calculated separately for beneficiaries participating in FP7 or Horizon 2020 EU grants for the first time and beneficiaries participating in the subsequent rounds. Assuming that more productive research institutes are located in the EU-15 countries and that collaboration with productive research partners from EU-15 increases the amount of funding, the grant amounts of EU-13 beneficiaries that collaborated in grants with EU-15 beneficiaries were also calculated. Similarly, the average grant amount of EU-15 beneficiaries that participated in grants without EU-13 beneficiaries was compared to the grant amount of the EU-15 beneficiaries that collaborated with at least one EU-13 beneficiary. For the descriptive statistical calculations the R software was used.^17^

### Regression analyses

First, univariate regression analyses were conducted to test the effect of country-level indicators on the total grant amounts assigned to the EU member states. The dependent variable was the total grant amount of the country (the sum of the funding assigned to all a country’s beneficiaries per country) per 100,000 inhabitants, for the entire period. We investigated the following four country-level indicators that potentially influence the grant distribution: (1) average gross domestic product per capita (GDP per capita) between 2007 and 2016, (2) average population size between 2007 and 2016, (3) disease burden in disability-adjusted life years (DALY) per 100,000 inhabitants in 2010, and (4) excellence of research capabilities, measured as the citation frequency of scientific literature published from the particular country between 1996 and 2016.

The average population size and GDP per capita values over the study period were calculated from annual data downloaded from the Eurostat database.^18^ The DALY values of 2010 were retrieved from the World Health Organisation (WHO) databank.^19^ The excellence of research capabilities was determined with the Scimago Journal & Country Rank database under the topic of “health professions”.^20^ The investigated parameter was the country-specific number of citations per document published between 1996 and 2016. The complete dataset of the explanatory variables by country can be found in Appendix 1.

The assumption of linearity was checked between the grant amount per 100,000 inhabitants and the explanatory variables with scatterplots (see Appendix 2). The scatterplot on GDP per capita showed that Luxembourg had a very high GDP per capita compared to the rest of the countries (see Appendix 2.1). This result had a negative effect on the positive correlation between GDP per capita and the grant amount per 100,000 inhabitants. According to a one-sided Dixon’s Q-test for outliers (Dixon test R-package), the GDP per capita in Luxemburg could be considered as an outlier (Q = 0.549; p<0.001). Nevertheless, we included Luxemburg in the analysis, but this should be taken into account when the results of the association between GDP per capita with the allocated funds are interpreted. Non-linearity was found for the population variable: countries with a high population had a similar grant amount per 100,000 inhabitants to countries with small populations (see Appendix 2.4). However, countries with a medium population size had a higher amount per 100,000 inhabitants. Because of this non-linearity, quartiles of the population size were tested as categorical variables in the regression analyses. The other three variables showed a clear linear relationship on the scatterplot. Therefore, standard linear regression analyses were used. T-tests were used to test the significance of the model coefficients. A level of 0.05 was considered to be significant throughout the analyses. We used STATA 15.0 to conduct the regression analyses.^21^

We conducted a multiple regression analysis to determine the effect of explanatory variables when adjusted for other factors. Those variables which did not influence the size of the effect of other factors were considered non-significant and excluded from the model. Regression diagnostics included checking normality with a normal quartile plot of the residuals, homoscedasticity with a residual versus fitted plot and multicollinearity with a calculation of the variance inflation factors (VIF) by explanatory variables and overall.^22^

## Results

### Overall grant distribution

A total of 5,810,052,343 EUR was distributed in the included healthcare research grants across the EU countries from 1 January 2007 to 31 December 2016. The average grant amount that was assigned to a country was 207,501,869 EUR with a median of 40,685,615 (IQR: 272,024,597). The number of unique participating beneficiaries (organizations that took part at least once in a grant during the included period) was 3,660, while the total number of beneficiaries was 12,678. Accordingly, an average beneficiary participated in 3.46 grants during the study period. The ten beneficiaries with the highest grant rates participated in 125.3 grants on average. All of these top ten beneficiaries were located in EU-15 countries.

The top five countries with the most total funding (the United Kingdom, Germany, the Netherlands, France and Italy) received 68.2% of the total EU healthcare grants. The five countries that had the most funding per 100,000 inhabitants (the Netherlands, Sweden, Denmark, Ireland and Belgium) were assigned with 48.7% of the total amount. The distribution of the total grant amounts among countries and the distribution of the grant amounts per 100,000 inhabitants are shown in Table 1. There was not any EU-13 country among the top five countries with the most total funding. Similar results were observed when funding was calculated for 100,000 inhabitants.

**Table 1:** Total received grant amounts between 2007 and 2016

| Country | FP7/H2020 health research grants per 100,000 inhabitants between 2007 and 2016 (in EUR) | Total FP7/H2020 health research grants between 2007 and 2016 (in EUR) |
| --- | --- | --- |
| Netherlands | 4,074,757 | 680,670,210 |
| Sweden | 3,352,475 | 318,401,349 |
| Denmark | 3,257,336 | 181,903,930 |
| Ireland | 2,782,080 | 127,457,470 |
| Belgium | 2,450,295 | 270,202,757 |
| Finland | 2,223,829 | 120,066,421 |
| United Kingdom | 1,824,941 | 1,158,140,380 |
| Austria | 1,794,676 | 151,556,376 |
| Luxembourg | 1,500,282 | 7,904,790 |
| Estonia | 1,236,893 | 16,394,550 |
| Germany | 1,227,201 | 997,083,187 |
| France | 1,004,105 | 656,085,172 |
| Spain | 791,564 | 367,646,205 |
| Italy | 787,988 | 470,631,443 |
| Cyprus | 776,587 | 6,483,100 |
| Slovenia | 762,467 | 15,624,008 |
| Greece | 714,193 | 78,606,387 |
| Portugal | 427,556 | 44,862,866 |
| Hungary | 367,203 | 36,508,364 |
| Croatia | 308,180 | 13,154,281 |
| Latvia | 232,236 | 4,793,065 |
| Czech Republic | 221,762 | 23,246,644 |
| Lithuania | 152,247 | 4,621,505 |
| Malta | 113,927 | 477,965 |
| Slovakia | 103,966 | 5,616,475 |
| Poland | 95,492 | 36,334,965 |
| Bulgaria | 62,414 | 4,576,343 |
| Romania | 54,627 | 11,002,147 |

The average grant amount assigned to a beneficiary was 458,278 EUR (median: 314,545; IQR: 411,537). Almost half of the distributed amount was concentrated around the 100 most successful beneficiaries. The top 100 beneficiaries with the most overall funding (2.73% of the total unique participating beneficiaries) had 45.8% of the total amount of the included grants. The participation history of beneficiaries showed a significant effect on the assigned grant amount. Beneficiaries who participated for the first time in EU healthcare grants during the analysed timeframe had an average grant amount of 367,093 EUR (median: 241,664; IQR: 358,602) per grant. Beneficiaries who previously participated in EU healthcare grants during the studied period were assigned with a larger average grant amount of 495,286 EUR (median: 349,619; IQR: 430,869) per grant. Thus, the average grant amount increased 34.9% when a beneficiary had previously participated in a grant (p < 10^−3^).

### Comparison of EU-15 with EU-13 countries

An overview of descriptive statistical calculations for EU-15 and EU-13 countries can be found in Table 2. From the total grant amount during the study period (5,810,052,343 EUR) 96.9% of the grants were assigned to EU-15 country beneficiaries. The number of unique EU-15 and EU-13 beneficiaries that participated in the FP7 or Horizon 2020 grants during the study period were 3,259 and 401, respectively. The total number of EU-15 beneficiaries who participated was 11,854 (1,446 coordinators and 10,408 participants), and the total number of EU-13 beneficiaries who participated was 824 (31 coordinators and 793 participants). Accordingly, the average number of grants were 3.6 and 2.1 for beneficiaries from EU-15 countries and from EU-13 countries, respectively.

**Table 2:** Overview of descriptive statistical calculations for EU-15 and EU-13 countries

|  | **EU-15** | **EU-13** | **Overall** |
| --- | --- | --- | --- |
| **Total grant amount** | 5,631,218,931 EUR<br>(96.9% of the overall) | 178,833,412 EUR<br>(3.1% of the overall) | 5,810,052,343 EUR |
| **Number of coordinating beneficiaries** | 1,446<br>(97.9% of the overall) | 31<br>(2.1% of the overall) | 1,477 |
| **Number of participating beneficiaries** | 10,408<br>(92.9% of the overall) | 793<br>(7.1% of the overall) | 11,201 |
| **Number of unique organizations as beneficiaries** | 3,259<br>(89.0% of the overall) | 401<br>(11.0% of the overall) | 3,660 |
| **Average participation per beneficiary between 2007-2016** | 3.6 | 2.1 | 3.5 |
| **Average grant amount per beneficiary** | 475,048 EUR<br>(median = 330,000) | 217,031 EUR<br>(median = 150,671) | 458,278 EUR<br>(median = 314,545) |
| **Average grant amount for first participation in the period** | 386,064 EUR<br>(median = 256,800) | 212,913 EUR<br>(median = 144,000) | 367,093 EUR<br>(median = 241,664) |
| **Average grant amount for subsequent participation in the period** | 508,788 EUR<br>(median = 359,928) | 220,934 EUR<br>(median = 171,520) | 495,286 EUR<br>(median = 349,619) |
| **Average grant amount for collaboration between EU-15 and EU-13 beneficiaries** | 411,590 EUR<br>(median = 289,709) | 215,732 EUR<br>(median = 152,520) | N/A |
| **Average grant amount for collaboration between only EU-15 or only EU-13 beneficiaries** | 515,317 EUR<br>(median = 356,723) | 278,662 EUR<br>( median: 50,000) | N/A |
| **Average grant amount for the beneficiaries that collaborated in grants >10 times** | 608,303 EUR<br>(median = 577,955) | No EU-13 beneficiaries collaborated >10 times. | N/A |

EU-15 and EU-13 beneficiaries were assigned with an average amount of 475,048 EUR per grant (median: 330,000; IQR: 422,786), and with an average amount of 217,031 EUR per grant (median: 150,671; IQR: 232,949), respectively. Larger grant amounts after the first grant participation was also apparent in a subgroup analysis of EU-15 countries: First time EU-15 beneficiaries received an average grant amount of 386,064 EUR (median: 256,800; IQR: 365,189), and when they participated in subsequent grants they received 508,788 EUR on average (median: 359,928; IQR: 437,988). The difference was a 31.8% increase, p-value: < 10^−3^. In contrast, grant amounts in EU-13 countries were similar across beneficiaries, regardless of whether they were a first time participant (average: 212,913 EUR; median: 144,000; IQR: 247,125) or a subsequent grant beneficiary (average: 220,934 EUR, median: 171,520; IQR: 214,047). The difference was not significant statistically (p = 0.6).

Participation of EU-13 beneficiaries occurred in only 464 of the 1,529 grants (31.4%). In the 1,065 grants where only EU-15 beneficiaries participated, the average grant amount per beneficiary was 515,317 EUR (median: 356,723; IQR: 443,080), while EU-15 beneficiaries had an average amount of 411,590 EUR (median: 289,709; IQR: 382,969) in grants when they collaborated with at least one EU-13 beneficiary. In these grants, EU-13 beneficiaries had an average amount of 215,732 EUR (median: 152,520; IQR: 227,230). When only EU-13 beneficiaries participated (17 grants), they had an average amount of 278,662 EUR (median: 50,000).

There were 134 collaborations where beneficiaries participated together in more than 10 grants and all of them were from EU-15 countries. These beneficiaries had a significantly larger average amount of 608,303 EUR (median: 577,955; IQR: 237,335).

### Univariate regression analysis

In the country-level regression analyses, GDP per capita showed a statistically significant positive association with amount of assigned funding. Every 1,000 additional EUR in GDP per capita was associated with an additional 49,680 EUR per 100,000 inhabitants difference (p < 0.001; R^2^ = 0.48). The excellence of healthcare research capabilities also showed a significant positive association of 139,046 EUR per 100,000 inhabitants per additional citation per document (p < 0.001); R^2^ = 0.64). Disease burden showed a statistically significant negative association of - 97,410 EUR per 100,000 inhabitants per 1,000 additional DALY per 100,000 inhabitants (p = 0.003; R^2^ = 0.30). There was no significant association between a country’s population size and the grant amount per 100,000 inhabitants (p = 0.3).

### Multiple regression analysis

In the initial multiple regression model (Appendix 3) only research excellence showed a significant association (p < 0.001). The GDP per capita showed a borderline non-significant association in the multiple regression model (GDP per capita: p = 0.097). The second and third quartiles of the population size showed a significant negative association compared to the first quartile (2^nd^ quartile: p = 0.031; 3^rd^ quartile: p = 0.048; 4^th^ quartile: p = 0.534; overall p = 0.082). The DALY per 100,000 inhabitants was not associated with the grant amount per 100,000 inhabitants (p = 0.222). This loss of significance was not unexpected, as GDP per capita was a confounding variable for the disease burden. The correlation plot in Appendix 4 shows that the countries with a lower GDP per capita have a higher disease burden (correlation = −0.7). Since disease burden showed a non-significant association on the grant amount per 100,000 inhabitants in the multiple analysis and because it was the least significant explanatory variable, DALY per 100,000 inhabitants was excluded from the final multiple regression model.

The final multiple regression model contained three significant explanatory variables that explained 82% of the variance of the allocated grants per 100,000 inhabitants. Results of the final multiple regression model is shown in Table 3. Research excellence adjusted for the GDP per capita and population size was significantly associated with the allocated grants (97,560 EUR per additional citation per publication; p < 0.001). GDP per capita adjusted for the population size and research excellence was also significantly associated (27,554 EUR per 1,000 additional GDP per capita; p = 0.002). Results on population size showed that countries with more inhabitants had more funding per 100,000 inhabitants than the smaller countries. The second and third quartiles differed statistically significantly from the first quartile, the forth quartile did not (p-values = 0.026, 0.024, 0.33).

**Table 3:** Results of the final multiple regression model

| Variable | Coefficient | P-value | Overall P-value | 95% Confidence interval | R<sup>2</sup> |
| --- | --- | --- | --- | --- | --- |
| **GDP per capita (1000 EUR)** | 27.55 | 0.002 | - | 11.32; 43.79 | 0.82 |
| **Research excellence** | 97,560 | <0.001 | - | 57,385; 137,736 |  |
| **Population (2nd quartile)** | 700,203 | 0.026 | 0.067 | 90,777; 1,309,630 |  |
| **Population (3rd quartile)** | 721,330 | 0.024 |  | 105,599; 1,337,061 |  |
| **Population (4th quartile)** | 293,956 | 0.328 |  | -315,950; 903,861 |  |

GDP: gross domestic product

No violations of the normality and homoscedasticity assumptions were detected. The VIF values for the remaining explanatory variables in the final multiple regression model were low with a mean VIF value of 1.48 for the final multiple regression model.

## Discussion

The distribution of healthcare grants among the included 28 EU countries showed a disproportionate distribution. Large countries such as the United Kingdom, Germany and France had more total amounts for healthcare research, but when the total grant amount was normalised for the population size, the Netherlands became the most successful country in PF7 and Horizon 2020 programs during the study period. This “Netherlands effect” proves that smaller countries can be competitive with larger countries. Our results are in line with conclusions of an earlier study, which confirmed that new EU Member States were generally under-represented for participation and coordination of health-related projects in the EU’s FP5 and FP6 programmes.^23^ A report on all FP7 projects (i.e. not only health-related) also found that EU-13 countries are benefitting less from their participation in FP7 than the EU-15 countries.^24^ A more recent report on participation found that in FP7 21% of all projects involved at least one EU-13 organizations, and in the Horizon 2020 programme this ratio had fallen to 17%.^25^

Several factors contributed to the large difference in total health-related grants in FP7 and Horizon 2020 between the EU-15 and the EU-13 countries (96.9% vs. 3.1%). The number of participating organizations were much less from EU-13 countries (3,259 vs. 401). Furthermore, beneficiaries from EU-15 countries were involved in more grants during the study period than from EU-13 countries (3.6 vs. 2.1). There was a significant difference in the average grant amounts between EU-15 and EU-13 beneficiaries (475,048 EUR vs. 217,031 EUR). The effect of subsequent participation by beneficiaries also contributed to unevenly distributed EU funding. The average grant amount for EU-15 beneficiaries rose by 31.8% compared to 3.8% for the EU-13 countries when they participated in at least one more project in the analysed period. Lastly, composition of the consortium also influenced the grant distribution, as grants with only EU-15 beneficiaries had a larger average amount per beneficiary than a mixture of EU-15 and EU-13 partners. Habitual couples of partners with collaboration in more than ten projects was observed only in EU-15 countries. Such beneficiaries had a larger average grant amount compared to those couples who collaborated in less than 10 projects. This finding reinforces results of an earlier study, which found that connecting to productive researchers who have a good collaboration network and flow of information increases the chances of securing more funding.^8^

The univariate regression analyses showed a significant positive association of GDP per capita and research excellence and a significant negative association of DALY with the grant amount per 100,000 inhabitants. These figures highlight that countries with a larger burden of disease had less funding for health-related research. Thus, the current allocation of EU funds may increase the health inequalities among countries. The positive correlation with research excellence showed that the countries with better research excellence were assigned more funding.

The results of the multiple regression analysis showed that when accounting for the effects of all potential factors at the same time, GDP per capita, research excellence and population size showed a significant association with the grant amount per 100,000 inhabitants, the latter having an inverted U-shaped relationship with it. As GDP per capita strongly correlates with disease burden, this correlation hides the observed univariate analysis finding on negative association of grant amounts with disease burden.

### Limitations

Our study has some limitations. The first limitation is the small sample size. Although the complete dataset contained 14,446 healthcare specific grants on the beneficiary level, the univariate and multiple analyses were executed on the aggregated country-level dataset (N = 28 EU Member States). This was done because the included explanatory variables were country-specific rather than beneficiary-specific. Second, the selected programmes in the Cordis database may not include all of the grants that are connected to healthcare research allocated from the EU. Grants on research topics such as nutrition and nanotechnology, which may be relevant in the field of healthcare were not included. Third, data from previous funding programmes such as the 6^th^ framework programme (FP6) were not included in this research. Lastly, comparisons were made between the beneficiaries that previously received EU funding and the beneficiaries that did not. However, we cannot exclude the possibility, based on the data presented here that the included beneficiaries may have received grants in previous funding programmes which could influence some findings.

## Conclusion

Our study showed a disproportionate country-group specific (EU-15 vs. EU-13) allocation of the total healthcare grants distributed in the FP7 and the Horizon 2020 programmes by the EU. Participation in previous grants and collaboration between strong research institutions increased the grant amount that was assigned per beneficiary, confirming the positive influence of a collaboration network in receiving research grants. EU grant allocation for healthcare research at a country-level seemed to depend on research excellence, which corresponds to our expectations. However, EU grant allocation for healthcare research also seemed to be dependent on the economic status and the size of the countries, while higher disease burden per 100,000 inhabitants apparently did not attract more EU research funding. As economic status is strongly correlated with the health status of the population the current funding mechanism further increases the gap in terms of health status between more and less developed countries.

In summary, our findings indicate that wealthier countries with a medium population size and strong medical research excellence had more EU funding for healthcare research than poor countries with a small or large population and less developed research in healthcare.

## Table legends

Table 1: Total assigned grant amounts between 2007 and 2016

Table 2: Overview of descriptive statistical calculations for EU-15 and EU-13 countries

Table 3: Results of the final multiple regression model

## Contributors

ZK raised the research idea and supervised the research. LHMA collected and cleaned the data, conducted the descriptive analyses and designed data visualisations with the support of MC and JGP. Regression analyses and the corresponding diagnostics were conducted by ZV. All authors contributed to research planning, analysis methodology development and manuscript preparation. The corresponding author attests that all listed authors meet authorship criteria and that no others meeting the criteria have been omitted.

